# Shotgun metagenomics of soil invertebrate communities reflects taxonomy, biomass and reference genome properties

**DOI:** 10.1101/2021.11.16.468383

**Authors:** Alexandra Schmidt, Clément Schneider, Peter Decker, Karin Hohberg, Jörg Römbke, Ricarda Lehmitz, Miklós Bálint

## Abstract

1. Metagenomics - shotgun sequencing of all DNA fragments from a community DNA extract - is routinely used to describe the composition, structure and function of microorganism communities. Advances in DNA sequencing and the availability of genome databases increasingly allow the use of shotgun metagenomics on eukaryotic communities. Metagenomics offers major advances in the recovery of biomass relationships in a sample, in comparison to taxonomic marker gene based approaches (metabarcoding). However, little is known about the factors which influence metagenomics data from eukaryotic communities, such as differences among organism groups, the properties of reference genomes and genome assemblies.
2. We evaluated how shotgun metagenomics records composition and biomass in artificial soil invertebrate communities. We generated mock communities of controlled biomass ratios from 28 species from all major soil mesofauna groups: mites, springtails, nematodes, tardigrades and potworms. We shotgun-sequenced these communities and taxonomically assigned them with a database of over 270 soil invertebrate genomes.
3. We recovered 90% of the species, and observed relatively high false positive detection rates. We found strong differences in reads assigned to different taxa, with some groups (e.g. springtails) consistently attracting more hits than others (e.g. enchytraeids). Original biomass could be predicted from read counts after considering these taxon-specific differences. Species with larger genomes, and with more complete assemblies consistently attracted more reads than species with smaller genomes. The GC content of the genome assemblies had no effect on the biomass-read relationships.
4. The results show considerable differences in taxon recovery and taxon specificity of biomass recovery from metagenomic sequence data. The properties of reference genomes and genome assemblies also influence biomass recovery, and they should be considered in metagenomic studies of eukaryotes. We provide a roadmap for investigating factors which influence metagenomics-based eukaryotic community reconstructions. Understanding these factors is timely as accessibility of DNA sequencing, and momentum for reference genomes projects show a future where the taxonomic assignment of DNA from any community sample becomes a reality.

## Introduction

Biodiversity research, and particularly the investigation of hard-to-observe ecological communities increasingly relies on DNA- and RNA-based tools (Taberlet, Bonin, Zinger, & Coissac, 2018). If preconditions are met, e.g. nucleotide sequence databases exist (Margaryan et al., 2021) with curated taxonomic links (Schenk, Hohberg, Helder, Ristau, & Traunspurger, 2017), and experimental designs are robust (Zinger et al., 2019), these approaches can provide much needed data on soil invertebrate diversity.

There are two main approaches to the molecular biomonitoring of communities: metabarcoding and metagenomics. Metabarcoding uses high-throughput-sequences of taxonomic marker genes (“barcodes”) which are PCR-amplified from a community DNA extract. Metabarcoding is becoming a standard tool in biodiversity research. Its use is supported by several years of research in distinct organisms groups (Taberlet et al., 2018), and the availability of barcode databases (Hebert, Cywinska, Ball, & deWaard, 2003; Nilsson et al., 2019). However, metabarcoding has an important long-known drawback: it relies on the amplification of a marker gene (Taberlet, Coissac, Pompanon, Brochmann, & Willerslev, 2012). This can result in biases in species recovery from the resulting sequence data, and in taxon-related distortions of the original biomass - sequencing read relationships (Piñol, Senar, & Symondson, 2019). However, the amplification step solves two important issues: one can effectively target the taxonomic groups of interest (e.g. insects), and avoid others (e.g. microorganisms) and small or rare organisms with low amounts of DNA can still be recorded. Metagenomics randomly sequences all DNA fragments from a community DNA extract, generally without enrichment of certain parts of the genome. It is more quantitative than metabarcoding, since it skips the potentially biased PCR amplification step of taxonomic marker genes (Bista et al., 2018). A random selection of DNA fragments is sequenced from the DNA extracts, resulting in a less biased representation of the community in the sequence data. The taxonomic assignment of metagenomic sequences needs genome databases, and consequently, metagenomics is more frequently applied on microbial communities, where more complete genomic resources are available (Parks et al., 2020). There are several approaches to circumvent this limitation, from mitogenomes (Arribas et al., 2020) to shallow genome sequencing (Bohmann, Mirarab, Bafna, & Gilbert, 2020). As genome sequencing technologies mature, the generation of reference genomes for all eukaryotes receives increasing attention (Lewin et al., 2018). However, the technical issues affecting metagenomics are much less investigated than issues affecting metabarcoding, at least for eukaryotes.

Here we evaluate how well metagenomics reflects composition and biomass in artificially composed (mock) communities of soil invertebrates. We use a large collection of soil invertebrate genomes to taxonomically assign metagenomic reads. We investigate the effects of metagenomic classification thresholds on correct and false identification. We evaluate the relationship between biomass and reads, and how this relationship is influenced by taxonomy and by the properties of the genome assemblies used for taxonomic assignments.

## Material & Methods

### Mock community construction

We constructed mock communities from 28 soil invertebrate species from six major taxonomic groups at the Senckenberg Museum of Natural History Görlitz. Specimens were either freshly collected and stored in 96% undenatured ethanol (Collembola, Gamasida, Oribatida), or they came from breeding cultures (Enchytraeidae, Nematoda, Tardigrada). Four different mock types were designed (Table 1). We varied the total body volume (the sum of body volumes of all individuals) across the four mock communities. The mocks contained very small species (Nematoda) with average body volumes per species of 0.10-0.15 × 10^-6^ μm^3^, up to large species (Collembola, Enchytraeidae, Gamasina, Oribatida) with average body volumes per species of 44.1-50.8 × 10^-6^ μm3 (Table 1). We used body volume as a proxy of biomass, and refer to it as biomass throughout the text. In the first mock, all species were represented with equal biomass. In the second mock, very small to small species had more biomass (200-500%) compared to medium and large species. In the third mock a part of very small to small species (7 of 11) had larger biomass (200-400%) than medium to large species. In the fourth mock most small species had more biomass than large species, but some medium to large species also had high biomass. All four mock types were replicated three times.

**Table 1.** Composition of mock communities. For species where different developmental stages were available, individuals of different sizes were used to achieve the necessary biomass [adults + juveniles, e.g. *Paramacrobiotus richtersi* in mock 1: 4 + 1].

|  |  |  | Number of individuals |  |  |  |
| --- | --- | --- | --- | --- | --- | --- |
| Taxon | mean body length [μm] | body volume [10<sup>-6</sup> μm<sup>3</sup>] | mock 1 | mock 2 | mock 3 | mock 4 |
| Tardigrada |  |  |  |  |  |  |
| *Paramacrobiotus richtersi* (Murray, 1911) | 700 | 12.1 | 4+1 | 9 | 0+9 | 2+5 |
| Nematoda |  |  |  |  |  |  |
| *Acrobeloides nanus* (de Man, 1880) | 340 | 0.15 | 355 | 1775 | 1420 | 710 |
| *Panagrolaimus detritophagus* Fuchs, 1930 | 380 | 0.10 | 521 | 1562 | 1562 | 521 |
| *Panagrellus redivivus* (Linnaeus, 1767) | 620 | 0.28 | 190 | 570 | 380 | 190 |
| *Poikilolaimus oxycerca* (de Man, 1895) | 930 | 0.98 | 54 | 162 | 54 | 162 |
| Collembola |  |  |  |  |  |  |
| *Sphaeridia pumilis* (Krausbauer, 1898) | 300 | 5.7 | 9 | 9 | 37 | 9 |
| *Proisotoma minuta* (Tullberg, 1871) | 880 | 11.0 | 5 | 4 | 5 | 5 |
| *Podura aquatica* Linnaeus, 1758 | 560 | 13.9 | 4 | 12 | 8 | 8 |
| *Desoria trispinata* (MacGillivray, 1896) | 1090 | 17.3 | 3 | 6 | 6 | 6 |
| *Isotomurus plumosus* Bagnall, 1940 | 1250 | 31.0 | 2 | 2 | 2 | 2 |
| *Deuterosminthurus bicinctus* (Koch, 1840) | 730 | 36.1 | 1+1 | 1+1 | 1+1 | 1+1 |
| *Sinella curviseta* Brook, 1882 | 1090 | 44.1 | 1 | 1 | 4 | 4 |
| *Folsomia fimetaria* (Linnaeus, 1758) | 1400 | 53.2 | 1 | 1 | 2 | 3 |
| Oribatida |  |  |  |  |  |  |
| *Tectocepheus velatus* (Michael, 1880) | 240 | 4.8 | 11 | 33 | 11 | 22 |
| *Minunthozetes semirufus* (C. L. Koch, 1841) | 280 | 5.6 | 9 | 28 | 19 | 10 |
| *Pantelozetes paolii* (Oudemans, 1913) | 340 | 12.9 | 4 | 12 | 4 | 8 |
| *Zygoribatula exilis* (Nicolet, 1855) | 360 | 13.7 | 4 | 12 | 8 | 12 |
| *Chamobates voigtsi* (Oudemans, 1902) | 300 | 15.9 | 3 | 3 | 7 | 3 |
| *Atropacarus striculus* (C. L. Koch, 1835) | 440 | 27.1 | 2 | 2 | 2 | 2 |
| *Liebstadia similis* (Michael, 1888) | 470 | 35.5 | 2 | 1 | 5 | 3 |
| *Eupelops occultus* (C. L. Koch, 1835) | 410 | 46.5 | 1 | 1 | 1 | 3 |
| *Oribatella quadricornuta* (Michael, 1880) | 560 | 50.8 | 1 | 1 | 2 | 2 |
| Gamasida |  |  |  |  |  |  |
| *Gaeolaelaps aculeifer* (Canestrini, 1883) | 700 | 22.0 | 2+1 | 5 | 2+1 | 5+6 |
| Enchytraeidae |  |  |  |  |  |  |
| *Enchytraeus bulbosus* Nielsen & Christensen, 1963 | 4000 |  | fragments |  |  |  |
| *Enchytraeus albidus* Henle, 1837 | 2500 |  |  |  |  |  |
| *Enchytraeus luxuriosus* Schmelz & Collado, 1999 | 10500 |  |  |  |  |  |
| *Enchytraeus bigeminus* Nielsen & Christensen, 1963 | 6500 |  |  |  |  |  |
| *Enchytraeus crypticus* Westheide & Graefe, 1992 | 7500 |  |  |  |  |  |

We used different formulas for body volume approximation. For Collembola we estimated body volumes as ellipsoid volumes (V(μm^3^) = 1.33 × π × a × b × c × 10^-6^, where a, b, c are axis lengths in μm). For Oribatida, Gamasida, and Enchytraeidae we estimated body volumes as cylinder volumes (V(μm 3) = π × L × r^2^ × 10^-6^, where L is height and r is radius), for Tardigrada V(μm^3^) = L × d^2^ × 0.785 × 10^-6^ (Hallas & Yeates, 1972), and for Nematoda V(μm3) = L × d2 × 0.577 × 10^-6^ was used (Andrássy, 1956). Average sizes of mite, nematode, tardigrade and springtail specimens were measured in the populations used for mock community construction, as body sizes can vary among specimens depending on life stage and other factors.

We used the tardigrade culture *Paramacrobiotus richtersi* (Murray, 1911) strain Hohberg-99 and the following cultures of nematodes: *Acrobeloides nanus* (de Man, 1880) strain Hohberg-99, *Panagrolaimus detritophagus* Fuchs, 1930 strain Hohberg-07, *Panagrellus redivivus* (Linnaeus, 1767) strain König-18 and *Poikilolaimus oxycerca* (de Man, 1895) strain Hohberg-01. Thousands of nematode specimens were extracted through sieves and milk filters from the culture plates into tap water. Nematode numbers and mean body volumes within the four stock solutions were then calculated by counting individuals of aliquots and measuring body length and width of 20 specimens per aliquot. After counting, we evaporated the water from each stock solution and added 96% ethanol. For the mock communities we added a calculated part of each of the stock solutions, holding the respective nematode volume, i.e. number × mean body volume. As enchytraeids are large compared to the other invertebrates, we used only body fragments. Tardigrades, collembolans and mites were individually counted into the mock communities. In order to achieve the needed biomass of the respective mock type, differently sized individuals (adults and juveniles) were used. All mock community samples were stored in 2 ml Eppendorf tubes in 96% undenatured ethanol at −20 °C until sequencing.

### Laboratory work and sequencing

Before performing the DNA extraction, ethanol was evaporated in a SpeedVac Concentrator Plus (Eppendorf, Hamburg, Germany) to avoid losing material. This is especially important for potentially floating Nematoda and Tardigrada specimens. DNA was extracted with DNeasy Blood and Tissue kit (Qiagen, Hilden, Germany). We included a negative control into the extractions to investigate possible cross-sample contamination. DNA concentration was measured on NanoDrop (Thermo Fisher Scientific, Waltham MA, USA) and Qubit^™^ (with the dsDNA BR Assay Kit (Thermo Fisher Scientific, Waltham MA, USA). Fragment length was checked on TapeStation 2200 (Agilent Technologies, Santa Clara CA, USA). Libraries were prepared with the NEB Next^®^ Ultra^™^ DNA Library Prep Kit (New England Biolabs, Ipswich MA, USA) and sequenced on an Illumina NovoSeq 6000 PE150 platform at Novogene (Hong Kong, China). Sequencing depth was 20 gigabase per mock community, and 1 gigabase for the negative control (2 × 150 bp, paired-end).

### Bioinformatics & data processing

Sequences were trimmed and quality checked with Autotrim v0.6.1 (Waldvogel et al., 2018). Autotrim relies on Trimmomatic (Bolger, Lohse, & Usadel, 2014), FastQC (Andrews, 2017/2021) and MultiQC (Ewels, Magnusson, Lundin, & Käller, 2016). It removes Illumina sequencing adapters, performs a quality control of the reads, and combines all information into a single report. Taxonomic classification was performed with Kraken2 v2.0.8 (Wood, Lu, & Langmead, 2019) against a designated soil invertebrate genome database (GenBank Bioproject PRJNA758215). This database contains short-read assemblies of over 250 species (FigShare doi: 10.6084/m9.figshare.16922890, Supplemental Table 1), including all species used for the mock communities. Before conducting metagenomic classification, the reference genomes were used to build a Kraken2 database with the default k-mer size (k=35). Taxonomic identification of reads was performed on 21 classification thresholds (between 0.0 to 1.0, at 0.05 increments). At each classification threshold, we accounted for possible contamination by extracting the hits of each taxon found in the negative control from the hits of that taxon in every mock community. We plotted correctly identified taxa, false negatives and false positives against the Kraken2 classification threshold, and selected the best-performing assignments for further analysis.

### Data analysis

Data analysis was conducted with R v3.6.1 in RStudio (Version 1.2.1335), with data formatted with tidyverse (Wickham et al., 2019). Graphs and plots were generated by using the package ggplot2 (Wickham, 2016). Unclassified reads, and classified reads representing less than 0.01 percent of the sample were removed from data. We evaluated false negatives and false positives at all 21 Kraken2 classification thresholds (FigShare doi: 10.6084/m9.figshare. 16922890).

We predicted read abundances with the total number of sequences obtained for each mock library with a generalized linear model. Initial independent variables were sequencing success, taxon group (Collembola, Enchytraeidae, Nematoda, Oribatida, Gamasida, Tardigrada), mock species biomasses, genome completeness (measured recovered complete Benchmarking Universal Single-Copy Orthologs, complete BUSCOs (Simão, Waterhouse, loannidis, Kriventseva, & Zdobnov, 2015)), GC content and genome sizes. We estimated genome sizes with ModEst, a new method which performs very well in comparison with flow cytometry measurements (Pfenninger, Schönnenbeck, & Schell, 2021). First we performed a combinatorial model selection with MuMIn (Burnham & Anderson, 2003). The best performing model based of quasi-AIC scores can be written up as hits ~ biomass + taxon_group + missing_buscos + genome_size. The final model was fitted with quasipoisson distribution to account for overdispersion. All predictors were scaled. Genome sizes were log-normalized before scaling. We evaluated the relative importance of the predictors by calculating model-specific variable importance scores in the R package vip (Greenwell & Boehmke, 2020).

We evaluated the correspondence between community composition captured by metagenomic reads and original biomass composition with redundancy analyses in vegan (Oksanen et al., 2019). We tested metagenomic hit model statistical significance with an ANOVA-like permutation test for redundancy analysis (Legendre & Legendre, 2012).

## Results

The sequencing resulted in ~69 million paired-end reads on average per mock community replicate, with a standard deviation of ~1.5 million reads. Raw sequencing results are available on the European Nucleotide Archive (accession number: PRJEB45431). About ten million reads were recorded in the negative control. Of the reads passing quality filtering, ~95 million were assigned to taxa at a 0.95 classification threshold (Table 1). The number of correctly classified species remained stable across all classification thresholds (Fig. 1). We retained results at 0.95 as a trade-off for correct and false classifications. Of the 28 species from the mock community, 27 were correctly identified at most classification thresholds (Fig 1). However, the number of false positive classifications strongly decreased at more stringent thresholds, from 181 to 11. The number of false negative classifications remained low, stable and consistent - a single species (an oribatid mite: Atropacarus striculus) was missed at most classification thresholds. Missing this species was due to the stringency of the bioinformatic sequence processing: the species yielded very few sequencing reads which were then discarded during data filtering.

**Fig. 1.**
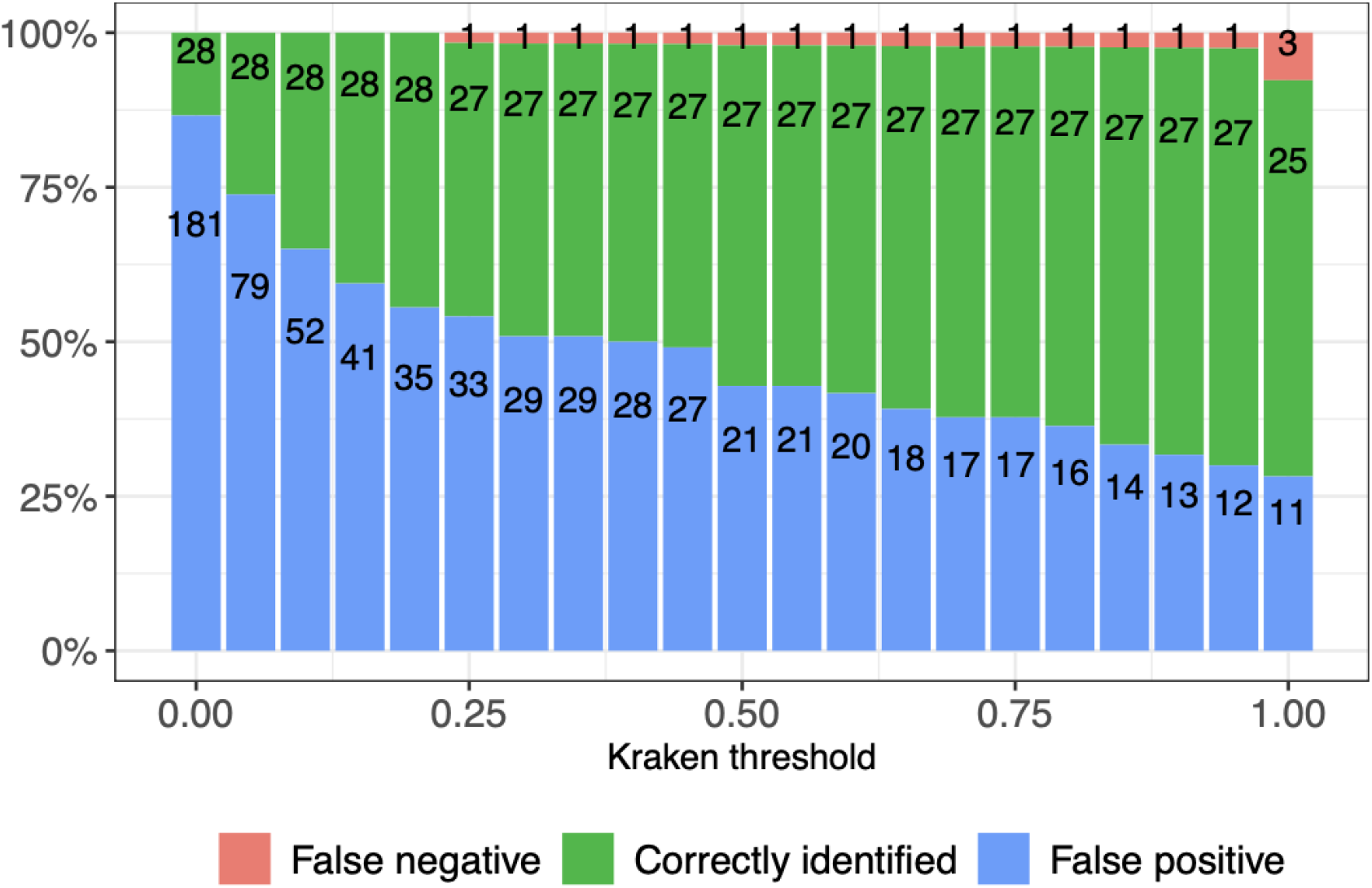
Species identification success along different Kraken2 classification thresholds. Numbers over bars represent the actual numbers of correctly identified species, and false negative and false positive identifications.

Some species consistently yielded more reads, regardless of their biomass ratios in the mocks (Fig. 2a). Sequencing depth differences among mock libraries and the GC content of the genomes had little predictive effect on assigned sequencing reads, so they were discarded during model selection. The final model (Fig. 2b, Table 2) showed that metagenomic sequencing success differed across the taxon groups. Compared to reads assigned to Collembola, assignment success to Tardigrada and Nematoda was slightly, but statistically insignificantly lower, while assignment success to Oribatida and Nematoda was statistically significantly lower (Table 2). Biomass of species was positively related to assigned metagenomic reads in all groups. Genome completeness had a statistically significant positive effect on metagenomic read assignment: overall more reads were assigned to taxa with more complete genomes, although this differed across taxon groups. Genome size had a statistically significant positive effect on metagenomic read assignment: more reads were assigned to taxa with larger genomes, regardless of the taxon group. Taxon groups were the most important predictors in the model (Fig. 2c). Replicates of the four mock community types were statistically significantly grouped together in the redundancy analysis (df = 3, F = 3.863, p < 0.001, Fig. 3).

**Fig. 2.**
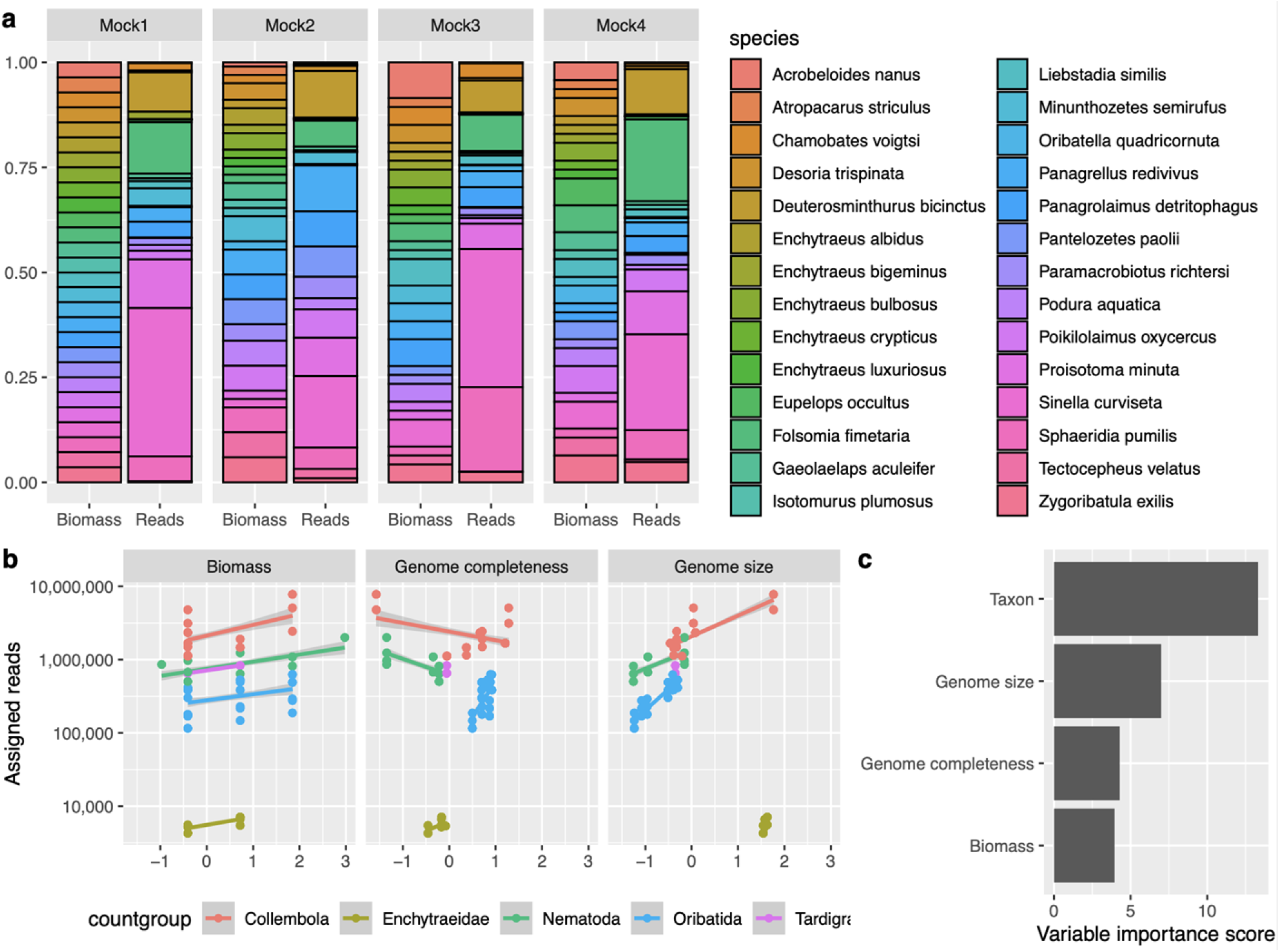
a) Biomass ratios of taxa and sequencing reads assigned to these taxa in four mock communities. b) GLM-predicted effects of biomass, genome completeness and genome size on taxonomically assigned metagenomic reads. c) Relative importance of GLM predictor variables.

**Table 2.** Model-predicted biomass, taxon group, genome completeness, genome size effects on assigned metagenomic read numbers. All predictors were scaled before model fitting. Genome size was log-normalized before scaling. Collembola served as a model intercept.

|  | Estimate | Standard error | t | *p* |
| --- | --- | --- | --- | --- |
| (Intercept) | 14.286 | 0.106 | 134.852 | 0.000 |
| Biomass | 0.215 | 0.055 | 3.942 | 0.000 |
| Enchytraeidae | -7.380 | 1.787 | -4.129 | 0.000 |
| Nematoda | 0.516 | 0.339 | 1.521 | 0.129 |
| Oribatida | -1.374 | 0.207 | -6.632 | 0.000 |
| Tardigrada | -0.372 | 0.356 | -1.043 | 0.298 |
| Genome completeness | 0.555 | 0.130 | 4.283 | 0.000 |
| Genome size | 1.165 | 0.167 | 6.991 | 0.000 |

**Fig. 3.**
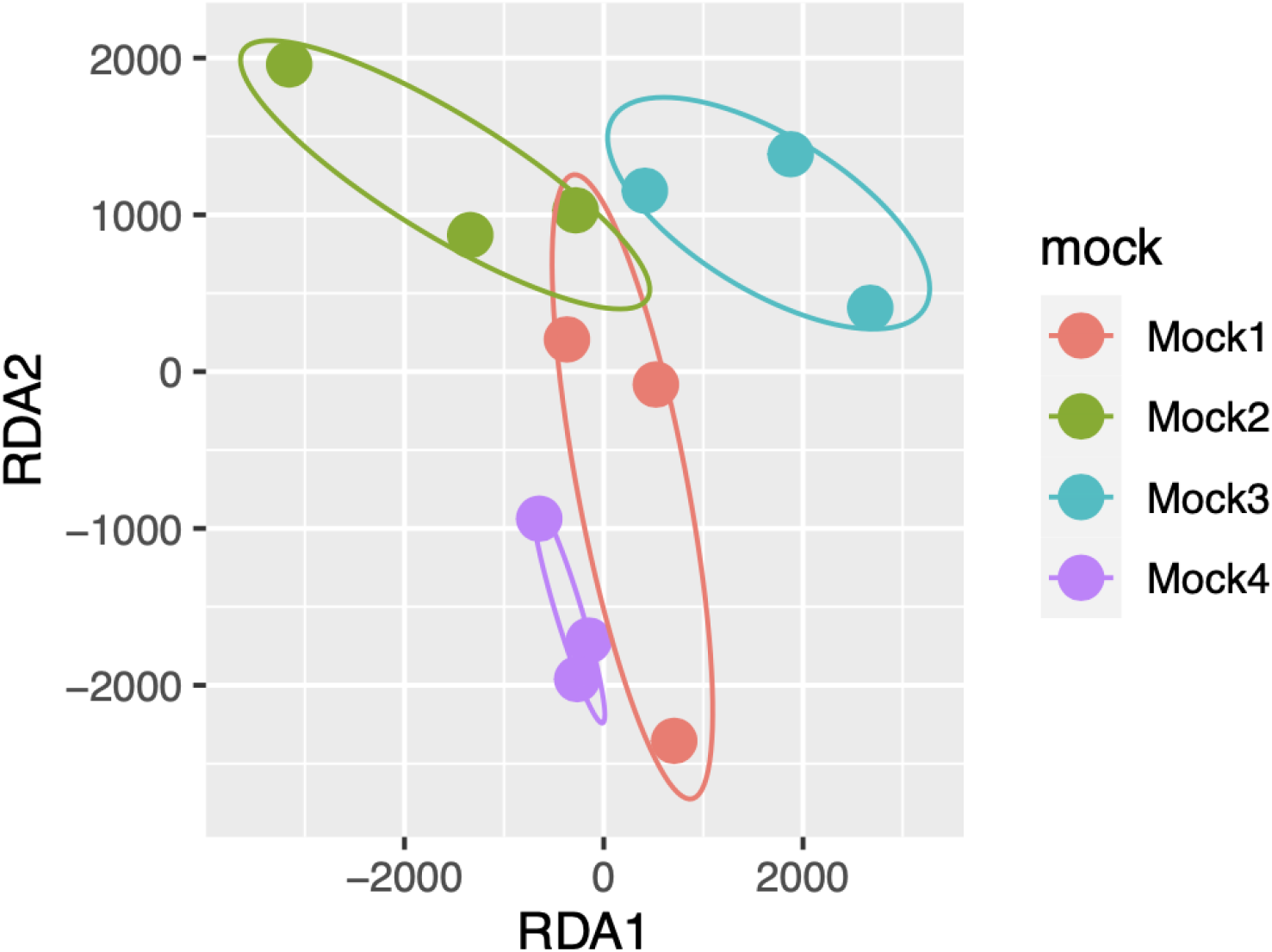
Redundancy analysis ordination of mock community replicates along the taxonomically assigned metagenomic reads.

## Discussion

We performed a shotgun metagenomic experiment on soil invertebrate mock communities of known composition. We assigned metagenomic reads to a genome database of soil invertebrates. We investigated how metagenomic reads record the presence of taxa in the mocks, whether read numbers reflect biomass, and how taxonomic, genome, and assembly properties influence biomass - read relationships.

Almost all species (27/28) were consistently detected at most classification thresholds. The single false negative species (*A. striculus*) was also detected with very low read numbers, and it was missed only because of stringent quality filtering. The number of false positives was high at low classification thresholds, and rapidly dropped at higher thresholds (Fig. 1). Eleven false positive assignments were retained even at the highest classification threshold. Possible explanations include contamination and bioinformatic issues. Cross-contamination is sometimes observed in mock metagenomes (Bista et al., 2018) but it cannot cause false positives here as all species were present in all mocks. Gut content may also result in the detection of unexpected taxa (Paula et al., 2016). However, most species used in these mocks are not predators. The predatory tardigrade *P. richtersi* was exclusively feeding on a nematode species which was also present in all the mock communities (*A. nanus*). The most likely explanation is related to some aspects of the metagenomic read assignment. The first candidate is the assignment algorithm itself, although comparisons show that Kraken is conservative (Harbert, 2018). Assignment of reads to closely related taxa is an unlikely cause since eight of the 12 false positive species (at 0.95 classification threshold) had no genus-level relatives in the mocks. Unmasked repeats might also erroneously attract reads during the assignment. Eukaryotes are rich in low complexity regions, and cross-assignment of these regions might be a considerable source of false positives in all eukaryotic metagenomes (Clarke et al., 2018). The effects of repeat regions in eukaryotic metagenomics assignments should be evaluated, although repeat identification is not trivial, especially for understudied taxa (Clarke et al., 2019).

The relationship between sequencing reads and the initial biomass of organisms is a central topic in the DNA-based analysis of community composition. In theory, more shotgun metagenomics reads should be assigned to species which are represented with higher biomass in a sample. However, this relationship might still be influenced by several factors. Here we investigated taxonomic effects, the impact of genome completeness, genome size, and GC content. We found that read counts were most strongly influenced by taxonomy, followed by genome size, genome completeness and biomass (Fig. 2c). We found no statistically significant effects of GC content on read assignment, although this was expected based on previous results with bacterial metagenomes (Browne et al., 2020).

There were consistently more reads assigned to some taxonomic groups than to others (Fig. 2b, Table 2). The impact of taxonomy on sequencing reads recovery seems to be systemic, with some species having many reads in all mocks, some species having only few reads (Fig. 2a), and one species was even missed due to the stringent filtering (Fig. 1). Species represented with low biomass in mocks were already found to result in false negatives in metagenomics (Bista 2018), and *A. striculus* was indeed represented with a relatively low biomass in the mocks. However, low biomass alone does not explain the strong taxon effect on read assignment. We suspect that the most important cause for the strong taxon effects is likely caused by differences in DNA yields among different taxa (Sato et al., 2019; Schiebelhut, Abboud, Daglio, Swift, & Dawson, 2017; Tourlousse et al., 2021). Some taxa, e.g. oribatid mites are very hardy, and their cuticles might present obstacles to tissue homogenisation during DNA extractions. Indeed, the single false negative species was an oribatid mite. Cells of different taxa might react differently to extraction (Costea et al., 2017; Morgan, Darling, & Eisen, 2010), with some species consistently yielding lower quality DNA in lower quantities (or no DNA at all) than others (Schiebelhut et al., 2017). However, differential DNA extraction efficiency does not explain why soft-bodied enchytraeids yield considerably less DNA than all other taxa (Fig. 2b). Differences in DNA content relative to body size (or biomass) might be responsible for this: some taxa may contain higher amounts of DNA per unit biomass than others. The association of DNA content with body size can be positive or negative depending on the organism group (Gregory, 2001).

Strong taxonomic effects on biomass-read relationships are interesting not only for metagenomic, but also for metabarcoding studies. It is generally assumed that primer mismatch is the most important source of taxonomically biased biomass-read relationships in metabarcoding (Collins et al., 2019; Lamb et al., 2019; Piñol et al., 2019). Our results suggest that taxon-specific differences in DNA extraction efficiency and/or DNA content might also play a role in taxonomic bias. However, recognizing this bias is difficult in metabarcoding: both primer bias, and factors influencing extraction DNA yields are likely phylogenetically conserved. Parallel metabarcoding and metagenomics studies on the same mock communities are necessary to evaluate the relative importance of primer bias versus DNA yield in biomass - read relationships (see e.g. (Bista et al., 2018).

Despite considerable taxonomic effects, biomass was a statistically significant predictor of reads (Fig. 2a, Table 2). This is in line with other metagenomic mock community studies on multicellular eukaryotes, such as benthic invertebrates (Bista et al., 2018) and pollen samples (Peel et al., 2019). The biomass effect on reads, although considerably smaller than taxon effects (Fig. 2c), was still sufficient to reflect compositional differences among the four mock types (Fig. 3). This confirms the suitability of shotgun metagenomics for a semi-quantitative comparison of soil invertebrate communities.

We found that reference genome properties influence taxonomic assignments and read-biomass relationships, and that these need to be considered in metagenomic studies on eukaryotes. We showed that reference genomes size influences metagenomic assignments, with larger genomes attracting more reads than smaller genomes (Fig. 2b). This is known from microbial studies where it was shown that average genome size of a microbial community influences metagenomics results (Beszteri, Temperton, Frickenhaus, & Giovannoni, 2010). We found that genome completeness recorded as BUSCO scores may also influence metagenomic assignments, with more complete genomes attracting more reads. This suggests that reference genome assembly properties should also be considered in metagenomic assignments, even though previous findings show that even low coverage reference genomes can perform well (Sarmashghi, Bohmann, P. Gilbert, Bafna, & Mirarab, 2019). GC content of genomes might also influence metagenomic assignments (Browne et al., 2020), although in our case this effect was limited (Table 2).

Our results outline a roadmap for future shotgun metagenomic work on metazoan mock communities. In the wet lab, DNA extraction needs to be optimized and likely adapted to taxa of interest. Differences in DNA content per unit biomass among and within major taxon groups should be evaluated and corrected for. In bioinformatics, assignment algorithms should be evaluated, adapted and developed with eukaryotes in mind. The performance of distinct genomic regions (i.e. conventional marker genes, mitogenomes, coding regions, ultraconserved regions, repeat elements) should be evaluated, especially with respect to false positive detections. Genome databases will likely remain incomplete for some time. An important direction is to evaluate how incomplete databases (i.e. databases not containing the target species, but congenerics or even less related species) perform in taxonomic assignments.

## Conclusion

Metagenomics is a promising alternative to m etabarcodi ng also for eukaryotic communities. Although theory suggests that metagenomic reads should well represent biomass relationships in communities, differences among organisms related to DNA extraction efficiency and genome properties have strong influences on the biomass - read relationships. These effects need to be further investigated and quantified in parallel metabarcoding - metagenomic experiments. The effects of taxonomy, genome and assembly properties should be considered in analyses. Generalized linear models provide an excellent opportunity for this. With affordable sequencing and increasingly accessible eukaryotic reference genomes metagenomics is becoming a viable alternative to metabarcoding for describing community composition and structure.

## Acknowledgements

The present study is a result of the LOEWE Centre for Translational Biodiversity Genomics (LOEWE-TBG) and it was funded through the programme “LOEWE – Landes-Offensive zur Entwicklung Wissenschaftlich-ökonomischer Exzellenz” of Hesse’s Ministry of Higher Education, Research, and the Arts. MB and CS acknowledge support from the German Research Foundation (DFG project BA 4843/4-1). For the verification of taxon material, we thank Axel Christian and Ulrich Burkhardt.

## Author contributions

MB, KH, RL conceived the ideas and designed methodology. RL, CS, KH, JR provided the animals and ensured correct taxonomic identification. AS, CS, PD, KH, JR, RL collected the data. AS processed the data. AS and MB analysed the data. MB led the writing of the manuscript. All authors contributed critically to the drafts and gave final approval for publication.

## Data availability

Sequence data is available in GenBank (PRJNA758215). R scripts and inputs are available in FigShare (doi: 10.6084/m9.figshare. 16922890).

## References

Andrássy, I. (1956). Rauminhalts- und Gewichtsbestimmung der Fadenwurmer (Nematoden). Acta Zoologica, 2, 1–15.

Andrews, S. (2021). S-andrews/FastQC [Java]. Retrieved from https://github.com/s-andrews/FastQC (Original work published 2017)

Arribas, P., Andújar, C., Moraza, M. L., Linard, B., Emerson, B. C., & Vogler, A. P. (2020). Mitochondrial Metagenomics Reveals the Ancient Origin and Phylodiversity of Soil Mites and Provides a Phylogeny of the Acari. Molecular Biology and Evolution, 37(3), 683–694. doi: 10.1093/molbev/msz255

Beszteri, B., Temperton, B., Frickenhaus, S., & Giovannoni, S. J. (2010). Average genome size: A potential source of bias in comparative metagenomics. The ISME Journal, 4(8), 1075–1077. doi: 10.1038/ismej.2010.29

Bista, I., Carvalho, G. R., Tang, M., Walsh, K., Zhou, X., Hajibabaei, M.,… Creer, S. (2018). Performance of amplicon and shotgun sequencing for accurate biomass estimation in invertebrate community samples. Molecular Ecology Resources, 18(5), 1020–1034. doi: https://doi.org/10.1111/1755-0998.12888

Bohmann, K., Mirarab, S., Bafna, V., & Gilbert, M. T. P. (2020). Beyond DNA barcoding: The unrealized potential of genome skim data in sample identification. Molecular Ecology, 29(14), 2521–2534. doi:https://doi.org/10.1111/mec.15507

Bolger, A. M., Lohse, M., & Usadel, B. (2014). Trimmomatic: A flexible trimmer for Illumina sequence data. Bioinformatics, 30(15), 2114–2120. doi: 10.1093/bioinformatics/btu170

Browne, P. D., Nielsen, T. K., Kot, W., Aggerholm, A., Gilbert, M. T. P., Puetz, L.,… Hansen, L. H. (2020). GC bias affects genomic and metagenomic reconstructions, underrepresenting GC-poor organisms. GigaScience, 9(2). doi: 10.1093/gigascience/giaa008

Burnham, K. P., & Anderson, D. R. (2003). Model Selection and Multimodel Inference: A Practical Information-Theoretic Approach. Springer Science & Business Media.

Clarke, E. L., Lauder, A. P., Hofstaedter, C. E., Hwang, Y., Fitzgerald, A. S., Imai, I.,… Collman, R. G. (2018). Microbial Lineages in Sarcoidosis. A Metagenomic Analysis Tailored for Low–Microbial Content Samples. American Journal of Respiratory and Critical Care Medicine, 197(2), 225–234. doi: 10.1164/rccm.201705-0891OC

Clarke, E. L., Taylor, L. J., Zhao, C., Connell, A., Lee, J.-J., Fett, B.,… Bittinger, K. (2019). Sunbeam: An extensible pipeline for analyzing metagenomic sequencing experiments. Microbiome, 7(1), 46. doi: 10.1186/s40168-019-0658-x

Collins, R. A., Bakker, J., Wangensteen, O. S., Soto, A. Z., Corrigan, L., Sims, D. W.,… Mariani, S. (2019). Non-specific amplification compromises environmental DNA metabarcoding with COI. Methods in Ecology and Evolution, 10(11), 1985–2001. doi: https://doi.org/10.1111/2041-210X.13276

Costea, P. I., Zeller, G., Sunagawa, S., Pelletier, E., Alberti, A., Levenez, F.,… Bork, P. (2017). Towards standards for human fecal sample processing in metagenomic studies. Nature Biotechnology, 35(11), 1069–1076. doi: 10.1038/nbt.3960

Ewels, P., Magnusson, M., Lundin, S., & Käller, M. (2016). MultiQC: Summarize analysis results for multiple tools and samples in a single report. Bioinformatics, 32(19), 3047–3048. doi: 10.1093/bioinformatics/btw354

Greenwell, B. M., & Boehmke, B. C. (2020). Variable Importance Plots—An Introduction to the vip Package. The R Journal, 12(1), 343. doi: 10.32614/RJ-2020-013

Gregory, T. R. (2001). Coincidence, coevolution, or causation? DNA content, cellsize, and the C-value enigma. Biological Reviews, 76(1), 65–101. doi: 10.1111/j.1469-185X.2000.tb00059.x

Hallas, T. E., & Yeates, G. W. (1972). Tardigrada of the soil and litter of a Danish beech forest. Pedobiologia. Retrieved from https://scholar.google.com/scholar_lookup?title=Tardigrada+of+the+soil+and+litter+of+a+Danish+beech+forest&author=Hallas%2C+T.E.&publication_year=1972

Harbert, R. S. (2018). Algorithms and strategies in short-read shotgun metagenomic reconstruction of plant communities. Applications in Plant Sciences, 6(3), e1034. doi: https://doi.org/10.1002/aps3.1034

Hebert, P. D. N., Cywinska, A., Ball, S. L., & deWaard, J. R. (2003). Biological identifications through DNA barcodes. Proceedings. Biological Sciences, 270(1512), 313–321. doi: 10.1098/rspb.2002.2218

Lamb, P. D., Hunter, E., Pinnegar, J. K., Creer, S., Davies, R. G., & Taylor, M. I. (2019). How quantitative is metabarcoding: A meta-analytical approach. Molecular Ecology, 28(2), 420–430. doi: https://doi.org/10.1111/mec.14920

Legendre, P., & Legendre, L. F. J. (2012). Numerical Ecology. Elsevier.

Lewin, H. A., Robinson, G. E., Kress, W. J., Baker, W. J., Coddington, J., Crandall, K. A.,… Zhang, G. (2018). Earth BioGenome Project: Sequencing life for the future of life. Proceedings of the National Academy of Sciences, 115(17), 4325–4333. doi: 10.1073/pnas.1720115115

Margaryan, A., Noer, C. L., Richter, S. R., Restrup, M. E., Bülow-Hansen, J. L., Leerhøi, F.,… Bohmann, K. (2021). Mitochondrial genomes of Danish vertebrate species generated for the national DNA reference database, DNAmark. Environmental DNA, 3(2), 472–480. doi: 10.1002/edn3.138

Morgan, J. L., Darling, A. E., & Eisen, J. A. (2010). Metagenomic Sequencing of an In Vitro-Simulated Microbial Community. PLOS ONE, 5(4), e10209. doi: 10.1371/journal.pone.0010209

Nilsson, R. H., Larsson, K.-H., Taylor, A. F. S., Bengtsson-Palme, J., Jeppesen, T. S., Schigel, D.,… Abarenkov, K. (2019). The UNITE database for molecular identification of fungi: Handling dark taxa and parallel taxonomic classifications. Nucleic Acids Research, 47(D1), D259–D264. doi: 10.1093/nar/gky1022

Oksanen, J., Blanchet, F. G., Friendly, M., Kindt, R., Legendre, P., McGlinn, D.,… Wagner, H. (2019). vegan: Community Ecology Package. Retrieved from https://CRAN.R-project.org/package=vegan

Parks, D. H., Chuvochina, M., Chaumeil, P.-A., Rinke, C., Mussig, A. J., & Hugenholtz, P. (2020). A complete domain-to-species taxonomy for Bacteria and Archaea. Nature Biotechnology, 38(9), 1079–1086. doi: 10.1038/s41587-020-0501-8

Paula, D. P., Linard, B., Crampton-Platt, A., Srivathsan, A., Timmermans, M. J. T. N., Sujii, E. R.,… Vogler, A. P. (2016). Uncovering Trophic Interactions in Arthropod Predators through DNA Shotgun-Sequencing of Gut Contents. PLOS ONE, 11(9), e0161841. doi: 10.1371/journal.pone.0161841

Peel, N., Dicks, L. V., Clark, M. D., Heavens, D., Percival-Alwyn, L., Cooper, C.,… Yu, D. W. (2019). Semi-quantitative characterisation of mixed pollen samples using MinION sequencing and Reverse Metagenomics (RevMet). Methods in Ecology and Evolution, 10(10), 1690–1701. doi: 10.1111/2041-210X.13265

Pfenninger, M., Schönnenbeck, P., & Schell, T. (2021). Precise estimation of genome size from NGS data (p.2021.05.18.444645). doi: 10.1101/2021.05.18.444645

Piñol, J., Senar, M. A., & Symondson, W. O. C. (2019). The choice of universal primers and the characteristics of the species mixture determine when DNA metabarcoding can be quantitative. Molecular Ecology, 28(2), 407–419. doi: 10.1111/mec.14776

Sarmashghi, S., Bohmann, K., P. Gilbert, M. T., Bafna, V., & Mirarab, S. (2019). Skmer: Assembly-free and alignment-free sample identification using genome skims. Genome Biology, 20(1), 34. doi: 10.1186/s13059-019-1632-4

Sato, M. P., Ogura, Y., Nakamura, K., Nishida, R., Gotoh, Y., Hayashi, M.,… Hayashi, T. (2019). Comparison of the sequencing bias of currently available library preparation kits for Illumina sequencing of bacterial genomes and metagenomes. DNA Research, 26(5), 391–398. doi: 10.1093/dnares/dsz017

Schenk, J., Hohberg, K., Helder, J., Ristau, K., & Traunspurger, W. (2017). The D3-D5 region of large subunit ribosomal DNA provides good resolution of German limnic and terrestrial nematode communities. Nematology, 19(7), 821–837. doi: 10.1163/15685411-00003089

Schiebelhut, L. M., Abboud, S. S., Daglio, L. E. G., Swift, H. F., & Dawson, M. N. (2017). A comparison of DNA extraction methods for high-throughput DNA analyses. Molecular Ecology Resources, 17(4), 721–729. doi: https://doi.org/10.1111/1755-0998.12620

Simão, F. A., Waterhouse, R. M., Ioannidis, P., Kriventseva, E. V., & Zdobnov, E. M. (2015). BUSCO:Assessing genome assembly and annotation completeness with single-copy orthologs. Bioinformatics, 31(19), 3210–3212. doi: 10.1093/bioinformatics/btv351

Taberlet, P., Bonin, A., Zinger, L., & Coissac, E. (2018). Environmental DNA: For Biodiversity Research and Monitoring. Oxford, New York: Oxford University Press.

Taberlet, P., Coissac, E., Pompanon, F., Brochmann, C., & Willerslev, E. (2012). Towards next-generation biodiversity assessment using DNA metabarcoding. Molecular Ecology, 21(8), 2045–2050. doi: 10.1111/j.1365-294X.2012.05470.x

Tourlousse, D. M., Narita, K., Miura, T., Sakamoto, M., Ohashi, A., Shiina, K.,… Terauchi, J. (2021). Validation and standardization of DNA extraction and library construction methods for metagenomics-based human fecal microbiome measurements. Microbiome, 9(1), 95. doi: 10.1186/s40168-021-01048-3

Waldvogel, A.-M., Wieser, A., Schell, T., Patel, S., Schmidt, H., Hankeln, T.,… Pfenninger, M. (2018). The genomic footprint of climate adaptation in Chironomus riparius. Molecular Ecology, 27(6), 1439–1456. doi: 10.1111/mec.14543

Wickham, H. (2016). ggplot2: Elegant Graphics for Data Analysis (2nd ed.). Springer International Publishing. doi: 10.1007/978-3-319-24277-4

Wickham, H., Averick, M., Bryan, J., Chang, W., McGowan, L., François, R.,… Yutani, H. (2019). Welcome to the Tidyverse. Journal of Open Source Software, 4(43), 1686. doi: 10.21105/joss.01686

Wood, D. E., Lu, J., & Langmead, B. (2019). Improved metagenomic analysis with Kraken 2. Genome Biology, 20(1), 257. doi: 10.1186/s13059-019-1891-0

Zinger, L., Bonin, A., Alsos, I. G., Bálint, M., Bik, H., Boyer, F.,… Taberlet, P. (2019). DNA metabarcoding—Need for robust experimental designs to draw sound ecological conclusions. Molecular Ecology, 28(8), 1857–1862. doi: 10.1111/mec.15060

